# Division of labour promotes the spread of information in colony emigrations by the ant *Temnothorax rugatulus*

**DOI:** 10.1101/791996

**Authors:** Gabriele Valentini, Naoki Masuda, Zachary Shaffer, Jake R. Hanson, Takao Sasaki, Sara Imari Walker, Theodore P. Pavlic, Stephen C. Pratt

## Abstract

The fitness of group-living animals often depends on how well members share information needed for collective decision-making. Theoretical studies have shown that collective choices can emerge in a homogeneous group of individuals following identical rules, but real animals show much evidence for heterogeneity in the degree and nature of their contribution to group decisions. In social insects, for example, the transmission and processing of information is influenced by a well-organized division of labour. Studies that accurately quantify how this behavioural heterogeneity affects the spread of information among group members are still lacking. In this paper, we look at nest choices during colony emigrations of the ant *Temnothorax rugatulus* and quantify the degree of behavioural heterogeneity of workers. Using methods from both machine learning and network analysis, we identify and characterize four behavioural castes of workers – primary, secondary, passive, and wandering – covering distinct roles in the spread of information during each emigration. This detailed characterization of the contribution of each worker can improve models of collective decision-making in this species and promises a deeper understanding of behavioural variation at the colony level.

## Introduction

Group-living animals must often act as integrated collectives in order to reach consensus on important decisions [1]. Choosing where to live [2], deciding among foraging patches [3,4], or suddenly changing the direction of group motion [5] are just a few examples of collective decisions. Despite the diversity in behavioural mechanisms that have evolved to address these and similar problems, there are many commonalities in the underlying strategies for processing the information needed to make a choice. For example, collectives pool information to mitigate the effect of uncertainty and increase decision accuracy [6], which requires the spread of information from informed individuals to uninformed ones [5,7].

Insights about how information spreads among group members can be obtained from the tools of network science [8,9]. In this approach, the group is reduced to a set of nodes, each representing a single animal, connected through edges that represent pairwise interactions. A challenge of this approach is to correctly identify each interaction and information transfer event during a given group decision. Sometimes interactions can be precisely observed, as in food transfer [10], social dominance [11,12], and physical contact [13]. When precise observation is not possible, physical proximity of a pair of animals is often used as a criterion of whether they are interacting [14]. However, proximity alone does not necessarily mean that an interaction occurred or, more importantly, that there was any transfer of information between the pair [15,16].

Behavioural heterogeneity among group members poses an additional challenge to studying the spread of information in collectives. Many theoretical models of collective decisions assume that group members all behave similarly to each other [17]. This assumption has shown how simple rules acted on by a mass of identical individuals can produce complex and functional group-level outcomes [18,19]. However, a full understanding of collective behaviour must account for well-known differences in the behaviour of members of real groups [20]. In recent years, acknowledgement of these differences has driven research efforts in animal personality and behavioural syndromes [21]. In eusocial insects, behavioural variation can be observed both across colonies and across workers of the same colony [22]. A long research tradition on division of labour has explored its basis in worker age, physiology, morphology, and experience; its degree of development in different species and contexts; its adaptive response to internal and external demands; and the ultimate forces contributing to its evolution [23–25]. However, the role of division of labour in collective decision making is less well understood [26].

In this study, we analyse behavioural heterogeneity in the context of collective nest-site choice by the rock ant *Temnothorax rugatulus*. Ants of this genus nest in pre-formed cavities such as rock crevices, and their ability to emigrate into nests with consistent suites of characteristics has made them a model system for understanding collective decision making. Emigrations are organized by a minority of active ants who search for candidate sites, assess them, and then recruit nestmates to promising finds [18,27,28]. The probability of recruitment initiation depends on site quality; hence, visitor numbers grow more rapidly at a better site, favouring its eventual selection. Active ants generally recruit one another using a behaviour called tandem running, in which a single nestmate is led from the current nest to the candidate new home. Once a quorum of ants has arrived at a site, they switch to the faster method of social transport, in which the passive majority of the colony is simply carried to the new site [18]. This quorum rule amplifies the effect of quality-dependent recruitment and ideally leads to all ants being carried to the same site before any competing site has reached the quorum.

Prior work on *Temnothorax* has shown a distinction between the active minority of ants that organizes the emigration and the colony majority that makes no obvious contribution to the decision [18,27– 29]. In addition, there is evidence for differences in the degree and consistency of participation by active ants, such that an ‘oligarchy’ of very active ants may play an outsized role over successive emigrations [27]. This oligarchy seems to be further organized into two subgroups, leaders of tandem runs and their followers, with levels of communication between these two groups higher than within each of them. However, these studies have focused only on a minority of key individuals; they do not provide a complete characterization of the contribution to the final decision of each worker and of their role in the necessary spreading and processing of information.

To provide a more complete account of behavioural variation, we recorded key behaviours of every ant within colonies choosing between two nests of different quality. We characterized the level of behavioural heterogeneity observed among workers and used machine-learning methods to group them into separate behavioural castes. We focused on tandem runs and transports, events in which information is clearly transferred between ants. Their conspicuousness allowed us to reconstruct complete networks of pairwise interactions throughout single emigrations as well as over the course of multiple emigrations by the same colony. Using tools from network science [30] and from information theory [31], we looked at how information spreads across the members of the colony and how this process correlates with the division of labour among individual workers as well as across different behavioural castes.

## Material and methods

All data and source code are available in [32].

### Experimental subjects

We used three colonies of *T. rugatulus* ants (ID numbers 6, 208, 3004), each with one queen and, respectively, 78, 81, and 33 workers. Colonies were collected in the Pinal Mountains near Globe, Arizona (N 33° 19.000’, W 110° 52.561’), during 2009. They were housed in nests composed of a balsa wood slat (50 mm × 75 mm) sandwiched between two glass slides. In the centre of the slat was a rectangular cavity (25 mm × 33 mm) to house the colony while the top slide had a 2 mm hole that served as an entrance. The nest was kept in a plastic box (110 mm × 110 mm) that was provided with a water tube and an agar-based diet replaced on a weekly basis [33]. Each ant received a unique pattern of four paint marks (one on the head, one on the thorax, and two on the abdomen) to allow individual identification during data collection.

### Experimental procedure

Colony emigrations were observed in a rectangular arena (37 cm × 65 cm) whose walls were coated with Fluon to prevent ants from leaving. We positioned two candidate new nests at one side of the arena and the old nest housing a colony at the opposite side with a distance of 50 cm between new and old nests. The candidate nests differed in their quality; the good nest was as described above, and the mediocre nest was identical except that it had a larger entrance (5.5 mm diameter). *Temnothorax* ants are known to strongly prefer smaller nest entrances [34,35]. The ants were induced to emigrate by removing the top slide of their nest. Each colony emigrated five times with a rest interval of two to five days between emigrations.

### Data collection

Experiments were video recorded using three cameras with 1k resolution. One camera gave a bird’s-eye view of the whole arena; the other two were positioned above each candidate nest and captured the nest interior at sufficiently high resolution to make the ants’ paint marks identifiable. We manually reviewed these recordings to compile a complete record of each ant’s arrivals, departures, and recruitment acts (see Table 1 for a complete list of recorded actions). For each action, we noted the time of occurrence, the identity of the ant and the nest at which the action occurred. We also recorded the origin and destination of every tandem run and transport, as determined from the recording of the whole arena.

**Table 1.** Ethogram of individual behaviours observed in video-recorded emigrations. Each row shows the symbol and definition of a particular action, along with the set of identifiers used to specify where it occurred.

| *Symbol* | *Definition* | *Location* |
| --- | --- | --- |
| *E* | Enter a candidate nest | {*good, mediocre*} |
| *L* | Leave a candidate nest | {*good, mediocre*} |
| *LB* | Begin leading a tandem run | {*old* → *good*, *old* → *mediocre*,<br>*mediocre* → *good*} |
| *FB* | Begin following a tandem run | {*old* → *good*, *old* → *mediocre*,<br>*mediocre* → *good*} |
| *TB* | Transport a brood item (*i.e.*, egg, larva, pupa) into a candidate nest | {*good, mediocre*} |
| *TU* | Transport an unknown object (*i.e.*, detritus for use as building material) into a candidate nest | {*good, mediocre*} |
| *TA* | Transport an adult ant into a candidate nest | {*good, mediocre*} |
| *BC* | Be carried into a candidate nest | {*good, mediocre*} |
| *TBO* | Transport a brood item out of a candidate nest | {*good, mediocre*} |
| *TAO* | Transport an adult ant out of a candidate nest | {*good, mediocre*} |
| *BCO* | Be carried out of a candidate nest | {*good, mediocre*} |

### Behavioural features of individual ants

For each individual we measured a set of features capturing behaviour important to the emigration. These included counts of transports and tandem runs, latencies to key events such as first visit to a nest, and durations of visits and transports (Table 2). As ants can perish or lose their paint marks between trials, each ant’s features were averaged over the number of trials in which she participated. For the times of first visit to a nest and of first transport event, we normalized values in the unit interval [0,1] where zero represents the beginning of the trial and one represents the completion of the emigration. Features that are conditioned on the occurrence of a transport event (e.g., time of first transport) are defined only for those ants transporting items in a trial and averaged only over trials satisfying the condition.

**Table 2.** List of measured features, their definitions, domain and whether they are defined for all ants or only for those ants involved in transport activity.

| *Feature* | *Definition* | *Domain* | *Always defined* |
| --- | --- | --- | --- |
| *Visits* | Average number of visits to any potential new nest (*i.e.*, good or mediocre nest) | $[0, \infty)$ | ✓ |
| *Visit duration* | Average duration of visits to any potential new nest | $[0, \infty)$ | ✓ |
| *Visits both* | Proportion of emigrations in which the ant visits both the good and the mediocre nest before transporting an item or be carried | $[0,1]$ | ✓ |
| *Total visits duration* | Cumulative duration of visits to any potential new nest before transporting an item | $[0, \infty)$ | ✗ |
| *Time of 1<sup>st</sup> visit* | Average proportion of the emigration at which the ant visits a potential new nest for the first time | $[0,1]$ | ✓ |
| *Tandem runs led* | Average number of tandem runs led including only runs from the old nest to either of the two potential nests and from the mediocre nest to the good nests | $[0, \infty)$ | ✓ |
| *Tandem runs followed* | Average number of tandem runs followed including only runs from the old nest to either of the two potential nests and from the mediocre nest to the good nests | $[0, \infty)$ | ✓ |
| *Reverse tandem runs led* | Average number of reverse tandem runs led including only runs from the either of the two potential nests to the old nest | $[0, \infty)$ | ✓ |
| *Reverse tandem runs followed* | Average number of reverse tandem runs followed including only runs from the either of the two potential nests to the old nest | $[0, \infty)$ | ✓ |
| *Items transported* | Average number of items (*i.e.*, adult ant, brood, dirt or unknown object) transported towards any nest | $[0, \infty)$ | ✓ |
| *Transport duration* | Average duration of a round-trip transport of any item | $[0, \infty)$ | ✗ |
| *Time of 1<sup>st</sup> transport* | Average proportion of the emigration at which the ant transports her first item | $[0,1]$ | ✗ |
| *Be carried* | Averaged number of times an ant is carried towards any nest | $[0, \infty)$ | ✓ |

### Measures of heterogeneity

To quantify the degree of heterogeneity in the distribution of a behavioural feature, we used both Lorenz curves and Gini coefficients [36]. A Lorenz curve is a probability plot that assesses how much the distribution of a feature across individuals varies from a hypothetical uniform distribution. In particular, it plots the cumulative portion of the total amount of a certain feature (*e.g.*, total number of tandem runs) against the cumulative portion of the population of ants, with ants ordered by increasing values of the feature. When a feature is equally distributed across the population, the corresponding Lorenz curve is a line with slope one (line of uniformity); the higher the degree of nonuniformity of the distribution, the larger the area between the line of uniformity and the Lorenz curve. The Gini coefficient is the ratio of this area to the total area under the line of uniformity. A Gini coefficient of one indicates maximal heterogeneity (e.g., one individual performs all of the actions measured by the feature) while a value of zero means that all individuals contribute equally to the feature).

### Clustering analysis

We used a two-level clustering analysis to group ants into distinct categories based on their behavioural features. At the higher level, we considered only behavioural features that are defined for all workers (i.e., excluding features defined only for transporters). At the lower level, we redefined the selection of behavioural features on the basis of the results at the higher level using either all features or only those associated with visits and be-carried events (some features have zero variance and cannot be used in the clustering analysis). Data from each colony were analysed separately from each other to prevent differences in feature distribution from affecting the results of the clustering. To explore the possibility of feature reduction, for each clustering level we computed the Pearson correlation coefficients between each pair of variables (function `cor` in package `stats` of R 3.4.3). We then performed a clustering analysis on standardized values of the features using the *k*-means algorithm (package `stats`, function `kmeans`, *k* = 2) and visualized the results using principal components analysis (package `stats`, function `prcomp`). No rotations were necessary, and all features with non-zero variance were retained due to their limited number (between four and eleven features) and good spread between their corresponding vectors. Finally, we analysed the distribution of features across clusters to characterize regularities in the roles played by ants in each category during an emigration.

### Construction of recruitment networks

We represented recruitment events in each emigration as a directed network. Recruitment in these ants lends itself to this approach because it consists of pairwise events in which one ant either leads or carries another ant to a candidate nest. To construct a network for a given trial, each ant participating in that trial was represented as a separate node. For each recruitment event, we add a directed edge from the node representing the recruiter (*i.e.*, the leader of a tandem run or the carrier of a transportee) to the node representing the recruit (*i.e.*, the follower or the transportee). We also labelled each edge by the type of recruitment event it represented: transport of an adult ant, forward tandem run, or reverse tandem run. The latter two are distinguished by their direction: forward tandem runs start at the original nest and end at either of the candidate new nests; they are usually seen early in the emigration before the start of transport [37]. Reverse tandem runs start at a candidate site and end at the original nest; they generally occur after transport has begun.

In addition to building a network for each emigration, we also constructed an aggregate network for each colony. The aggregate network was built similarly to those for individual trials, but it included edges for every recruitment event in all trials of a single colony. Aggregate networks provide us with a visualization of the long-term interaction patterns among ants that is less susceptible to the limited chance that ants have to encounter and possibly interact with each other during a single emigration.

### Core of aggregate recruitment networks

For each colony, we analysed its aggregate recruitment network to separate ants into two sets: 1) those belonging to the core of the network, and 2) those instead belonging to its periphery. The core is a subgroup of highly interlinked nodes representing ants that frequently interact with each other. There are several definitions of the network core [38]. The one we adopted is a particular type of *k*-core, with *k* = 1, where each node in the core has at least one directed edge to another node in the core [39,40]. We determined the core by removing all nodes that did not have outgoing edges (*i.e.*, ants with no active recruitment events) and all edges directed at any of the removed nodes (i.e., events where the removed ants were recruited by others). As this pruning can result in new nodes without outgoing edges, we repeated the procedure until no more nodes could be removed (*i.e.*, either the network is empty, or all remaining nodes have at least one outgoing edge).

### Measures of division of labour

We used an information-theoretic framework that measures both the degree of specialization of each *ant* and the degree at which *tasks* are performed by the same individuals [31]. Specialization of individual workers, *i.e.*, their tendency to perform certain tasks over others, can be quantified by the index *DOL*_*indiv*_ = *I*(*ant; task*)/*H*(*task*), while the degree to which tasks are performed by the same individuals is defined as *DOL*_*task*_ = *I*(*ant; task*)/*H*(*ant*). In these equations, function *I* is the mutual information between two variables and function *H* is the Shannon entropy of one variable. *DOL*_*indiv*_ = 1 indicates that given the identity of an ant we have complete knowledge of the task performed by that ant, whereas *DOL*_*indiv*_ = 0 indicates that tasks are performed randomly by ants and no association is present. *DOL*_*task*_ = 1 indicates that given a task we have complete knowledge of the identity of the ant performing that task whereas *DOL*_*task*_ = 0 indicates that ants randomly perform all tasks. An aggregate measure of division of labour is their geometric mean 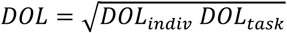. Values of *DOL* close to 1 correspond to a high degree of division of labour while values close to 0 represent instead the absence of any predictive relationship between a task to be performed and the ants that perform it. To compute these measures for each colony, we considered six tasks: leading a forward tandem run, following a forward tandem run, leading a reverse tandem run, following a reverse tandem run, transporting an item to a nest, and being carried to a nest. We then counted how many times each ant performed each task over the course of all emigrations. Using this data, we computed *DOL* measures both among the individuals of the colony (i.e., variable *ant* represents the unique id of each ant) and among behavioural castes (i.e., variable *ant* represents the role assigned to each ant during the clustering analysis). Mutual information and Shannon entropy were computed in R version 3.4.3 using the rinform package [41].

## Results

We observed a total of 15 emigrations in which a colony faced a choice between a *good* and a *mediocre* nest (five emigrations each by three colonies). The first trial by colony 208 was not analysed because a lighting disruption interfered with the quality difference between sites. In all but one emigration (colony 3004, trial 3), the colony made the expected choice and moved to the good nest. In two emigrations (colony 208, trials 2 and 3), the colony initiated parallel emigrations to both the good and the mediocre nest but later reunified at the good nest. From these 14 trials, we collected time-stamped behavioural data for each individual ant and analysed the contribution to the decision-making process of different members of the colony.

### Behavioural diversity of workers

As described previously for the related species *Temnothorax albipennis* [34], the contributions of individual workers in an emigration are far from equal. The majority of colony members experience the decision-making process only passively, when carried from one nest to another by a nestmate (Figure 1a, upper portion). A smaller portion of the workforce instead contributes actively to finding a new home and moving there (Figure 1a, lower portion). Initially, a few ants discover and perform short visits to the candidate nests (0–3.5 hours). Some of these ants later recruit other colony members through a number of tandem runs (3.5–4.5 hours). Finally, the bulk of the colony is carried to the chosen nest by this hard-working minority (4.5–6.5 hours).

**Figure 1.**
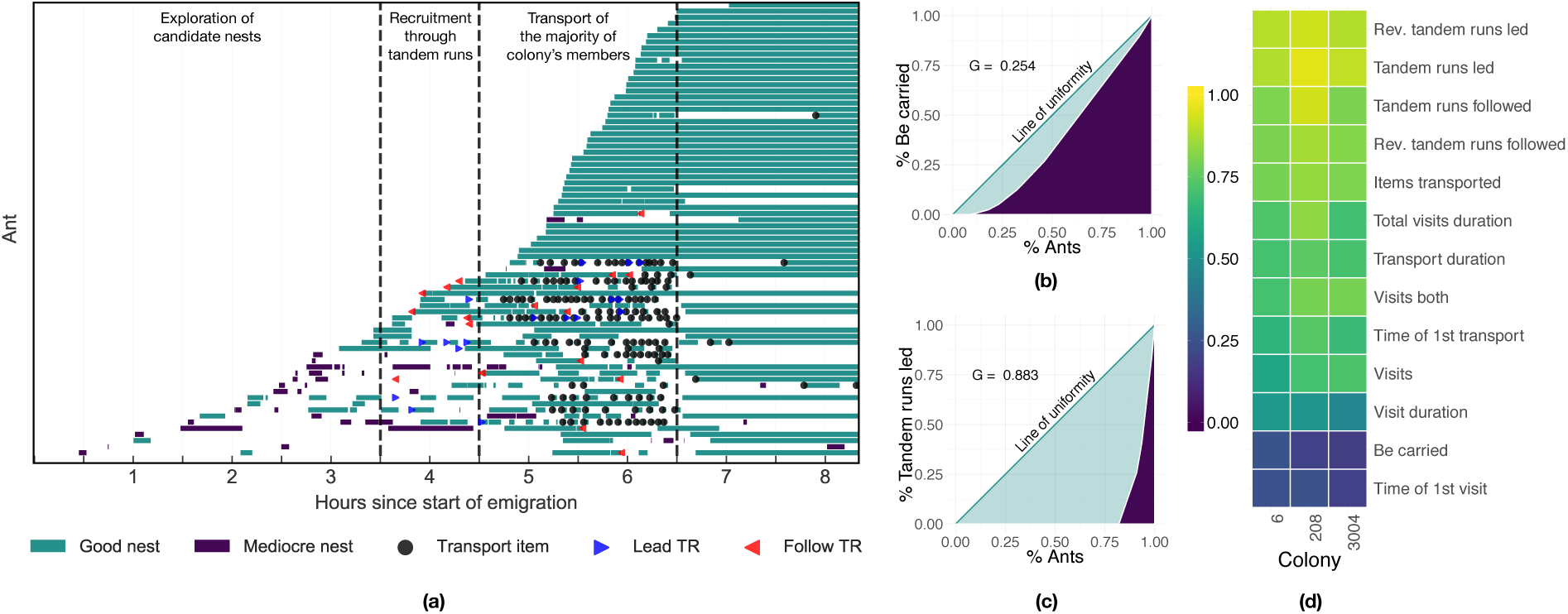
Panel (a) shows the individual behaviour over time of each ant in colony 6 during the first treatment. Horizontal bars denote time spent inside a candidate nest; symbols denote the occurrence of transport and tandem running behaviours; dashed vertical lines denote, from left to right, the exploration phase, the initial recruitment through tandem running, and the transport phase. Panel (b) and panel (c) show the Lorentz curves of the distribution of behaviours be-carried and leading tandem run across the members of colony 6 in all treatments. Panel (d) provides a heatmap representation of the Gini coefficients computed for a number of different behavioural features for each considered colony (higher values correspond to more unequal distributions).

For each colony, we measured the distribution of several behavioural features across colony members. Figure 1d provides a compact representation of how this distribution varied among behavioural features. Some behaviours were performed at similar frequency by all colony members. Being carried to a nest, for example, happens to most ants one time (Figure 1b), and the time of first visit was similar for most ants. Other behaviours, such as leading a tandem run, following a tandem run, or transporting a nestmate, had a much more unequal distribution, with only few workers performing them (Figure 1c). The remaining behavioural features, mostly concerned with visits to candidate nests, showed an intermediate level of diversity.

### Workers’ roles during the emigration

Even if only marginally, all ants in a colony behave differently from each other and therefore should be scored using a continuous scale. However, it is conceptually advantageous to separate them into a few distinct classes depending on their role in the collective decision. To do so, we first looked for correlations among behavioural features in each colony. Features related to tandem running and being carried generally have a low level of correlation with each other and with all other behavioural features (see Figure 2a for colony 6, Figure S1 for all colonies). In contrast, features related to the number and timing of visits and to the number of transported items are more correlated with each other. The numbers of visits and transported items are positively correlated with the likelihood of visiting both nests while the time of first visit is positively correlated with the average visit duration. The remaining correlations among this subset of features are all negative: ants that perform many visits, visit both nests, and/or transport many items make shorter visits and discover candidate nests earlier in the emigration process.

**Figure 2.**
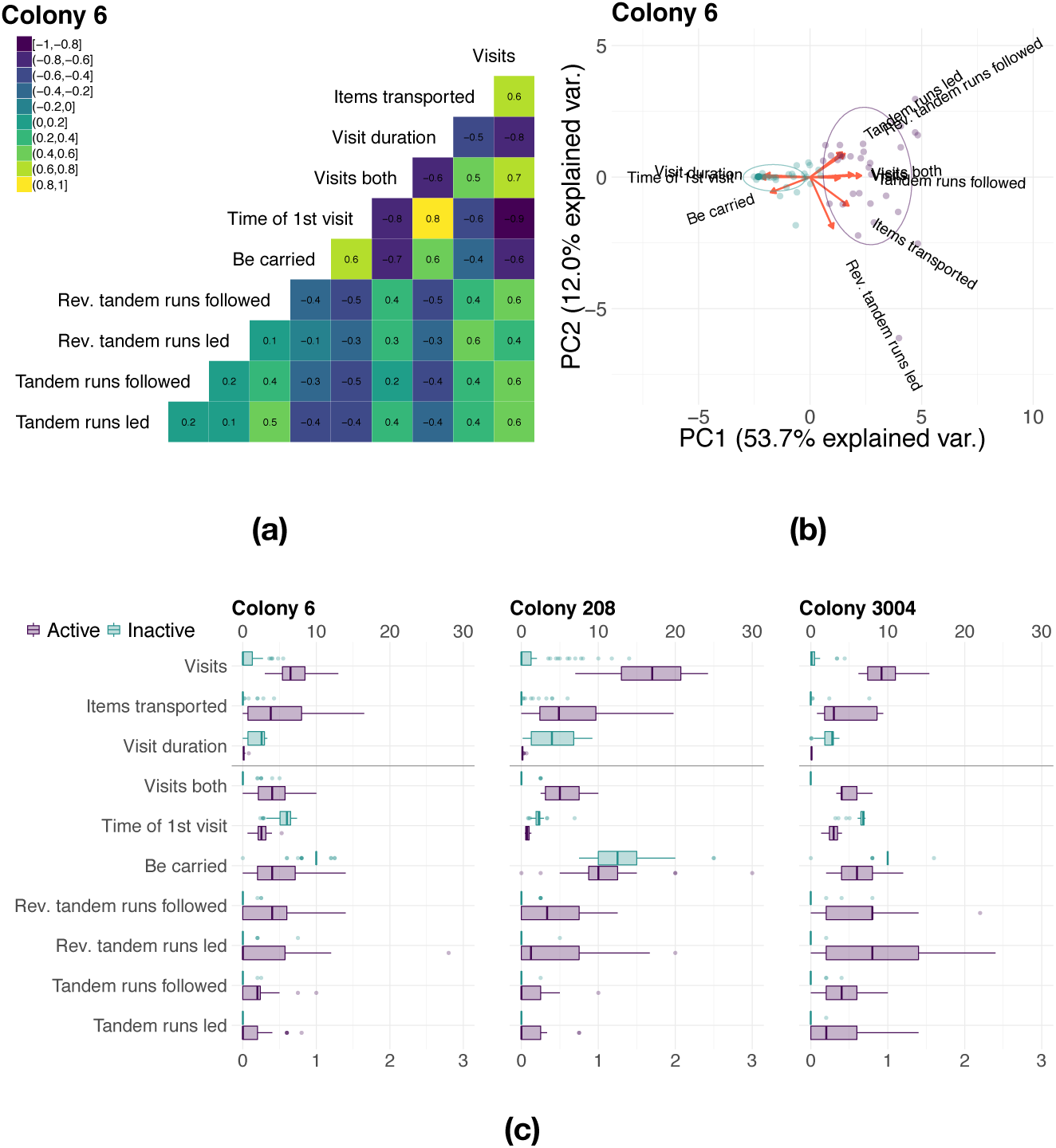
Illustration of the higher level of the clustering analysis. Panel (a) shows the matrix of Pearson correlation coefficients between behavioural features for colony 6. Panel (b) shows a PCA representation over the first two components of the PCA loadings and of clustered workers for colony 6 (purple symbols represent active ants; green symbols represent inactive ants). Panel (c) shows the distribution of behavioural features divided in active and inactive workers for all colonies (the average visit duration is reported in hours).

A principal component analysis (PCA) of the distribution of behavioural features (see Figure 2b for colony 6, Figure S2 for all colonies) shows that workers can be divided into two separate groups: ants that are carried to a candidate nest and ants that instead engage in transport and tandem running behaviours. When projected over the first two components of the PCA, which explain between 62% and 75% of the observed variance in each of the three studied colonies, the loadings form two sets approximately orthogonal to each other. On this basis, we performed a clustering analysis of each colony (*k*-means algorithm, data standardized) and looked for two clusters (see Figure 2c). As previously found for the case of task allocation in the same species [42], the clustering analysis separates *active* workers (*i.e.*, those that actively contribute to the emigration) from *inactive* ones (i.e., those that passively experience the emigration). Approximately a third of the workers of each colony (38%, 25%, and 27%, respectively, for colonies 6, 208, 3004) are classified as active ants.

The statistical classification described above lacks generalizability due to the limited number of trials. Inspection of the behavioural profiles of members of each category suggests more robust heuristic rules for classification. Active workers (see Figure 2c) are those that lead and follow most tandem runs, transport more items and therefore visit candidate nests more often; their visits are shorter, more likely to discover both nests, and happen earlier in the emigration process. Inactive workers are carried to a candidate nest approximately one time for each emigration, do not participate in tandem runs and do not transport items. They perform far fewer visits, generally visiting only one candidate nest (*i.e.*, the one where they have been carried to), and remain in that nest for longer than active workers. Based on these distributions of behavioural features, we defined as active workers any ant in the colony that 1) participates in at least one (forward or reverse) tandem run (as a leader or as a follower) or 2) transports at least one item. All workers that did not satisfy this definition were classified as inactive. After relabelling workers according to these criteria, roughly half the workers in each colony are active (50%, 40%, and 52% for colonies 6, 208, and 3004).

We next repeated the same clustering analysis for both active and inactive workers, to detect finer behavioural categories within each type. In both cases, behavioural features are generally characterized by a low level of pairwise correlation so that all considered features were retained and the results of PCA still show two separate groups of workers (see Section 1.2 of SI). We therefore clustered each higher-level group into two subgroups (Figure 3a and Figure S7): active ants are divided between *primary* workers and *secondary* workers; inactive ants are divided between *passive* workers and *wandering* workers.

**Figure 3.**
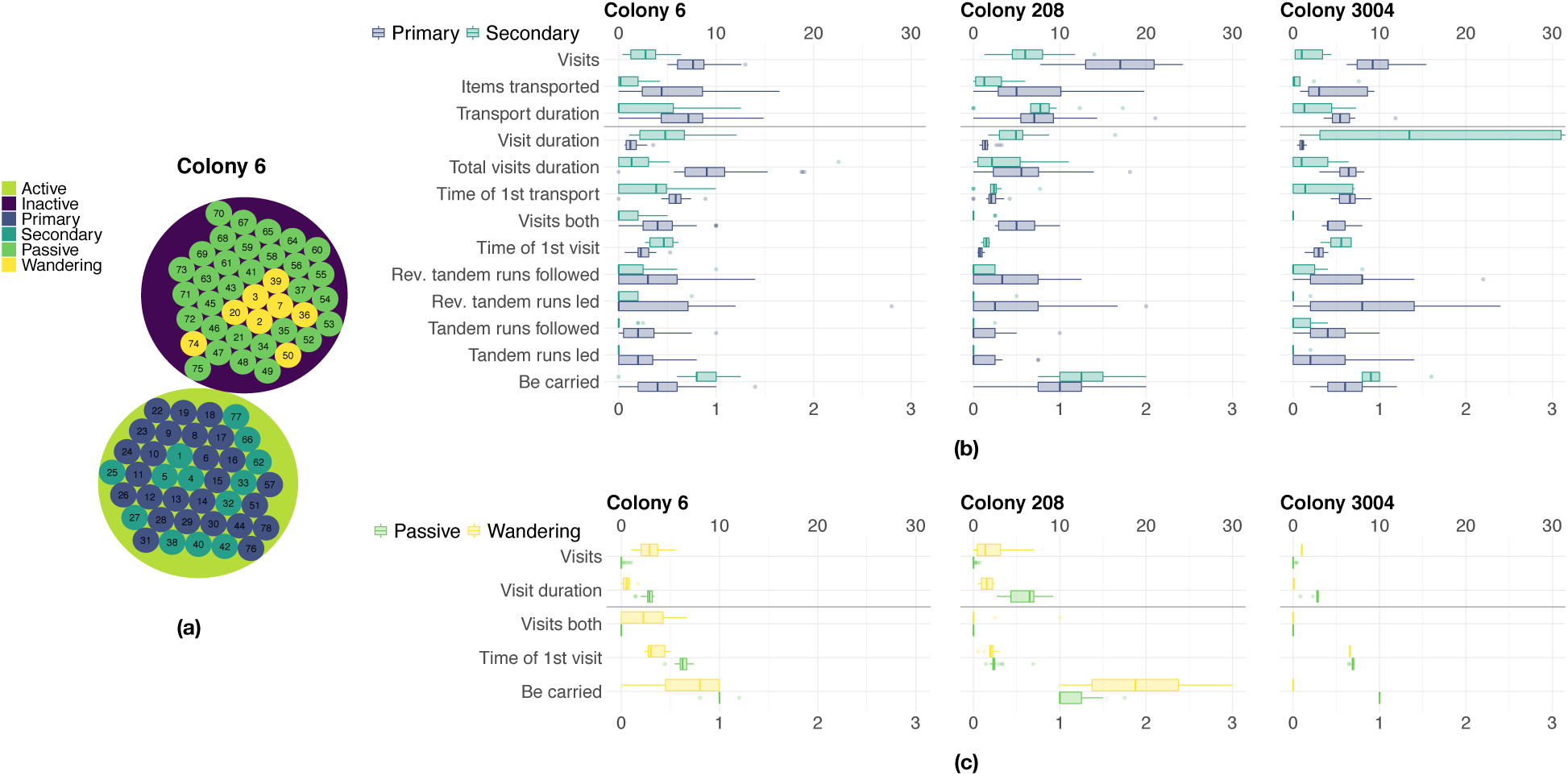
Illustration of the results of the clustering at the lower level of the hierarchy. Panel (a) shows the results of the two-level clustering of all ants for colony 6 as a circle plot. Panel (b) shows the results for active workers clustered in the groups primary and secondary ants. Panel (c) shows the results for the inactive workers clustered in the groups passive and wandering ants. The average visit duration and the total visit duration before transport are reported in hours; the average transport duration is reported in minutes.

Primary workers make up about a third of the colony (33%, 24%, and 27%, respectively, for colonies 6, 208, and 3004). These workers (cf. Figure 3b) visit candidate nests more often; their visits are generally shorter than those of secondary workers but, cumulatively, they spend more time within these nests before beginning to transport items. Consequently, they begin transporting later, at about three fourths of the emigration process. Despite this, they tend to transport a marginally higher number of items. Primary workers are those that participate more frequently in tandem runs and that are carried less frequently to a candidate nest. In contrast, secondary ants (17%, 16%, and 24% of each colony) are generally transported to a candidate nest but, soon after, begin frequent journeys to the old nest to transport items back to the new home. Unlike primary workers, they do not visit both candidate nests and they begin transporting items earlier in the emigration process.

Passive workers (40%, 48%, and 46% of the colony, respectively, for colonies 6, 208, and 3004) are truly inactive ants (Figure 3c); they are carried to a nest only once (slightly more for colony 208 due to two colony reunifications) during the entire emigration and spend the rest of their time there without visiting other nests. Wandering workers (10%, 12%, and 3% of each colony) are inactive with respect to the decision-making process but still show high levels of activity in terms of their number of visits to candidate nests. They make frequent and brief visits but never contribute to the emigration by engaging in recruitment behaviours. One possibility is that wandering ants largely ignore the ongoing emigration while exploring the environment in search for food or water.

### Recruitment networks

The different contributions of primary, secondary, passive, and wandering ants are evident in the visualizations of recruitment networks (Figures S8–S10). The prominent role of primary ants in the final choice of the colony is evidenced by their hub-like positions in the networks (Figures 4a and 4b). Likewise, secondary ants often appear as hubs but with fewer outgoing edges than those of primary ants. Passive ants largely appear as leaves in the network, generally subject of only one recruitment event, while wandering ants either appear as isolated nodes or as nodes subject to a higher number of recruitment events.

**Figure 4.**
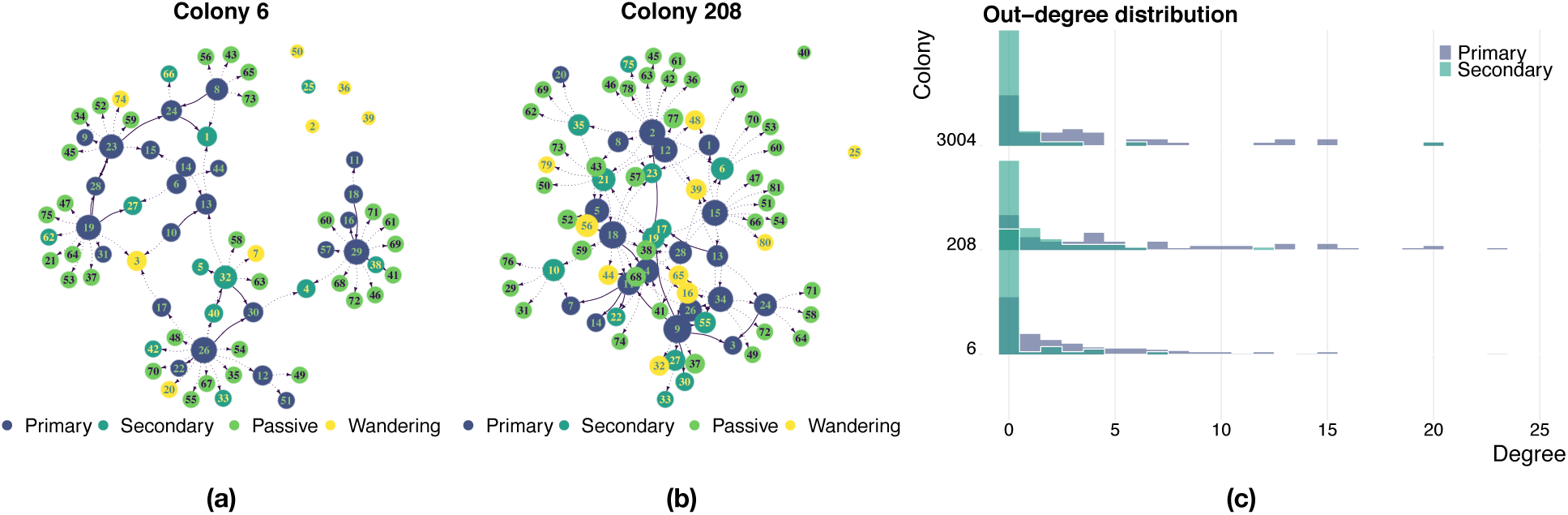
Illustration of recruitment networks and their outdegree distribution. Panel (a) shows the recruitment network for colony 6, treatment 1. Panel (b) shows the recruitment network for colony 208, treatment 2. Solid arrows represent tandem runs (both direct and reverse), dotted arrows represent transport events. Panel (c) shows the outdegree distribution for primary and secondary ants for all colonies.

Twelve of the 14 networks have a relatively simple structure composed of one or, sporadically, a few connected components and some isolated nodes (Figure 4a). The corresponding emigrations proceeded smoothly with most recruiters committed to the (eventually) chosen nest (eleven times the good nest, one time the mediocre nest) from early in the emigration; at most a few ants initially and only temporarily recruited to the alternative site. Two emigrations (colony 208, trials 2 and 3) differed from the rest in their markedly more complex recruitment networks, with only one large component and a higher edge density (see Figure 4b for colony 208, treatment 2). In these emigrations, the recruiters were initially split, with primary ants recruiting to both nests but eventually all choosing the better one. The higher edge density is therefore a result of this initial indecision and also of the additional recruitment effort necessary to later reunify the colony at a single site. Compared to the other emigrations, secondary ants in these two trials were more often involved in transport events and the corresponding hubs have a higher degree, albeit smaller than that of primary ants. A few passive ants were transported to both nests and therefore have two incoming edges. For most of them, as well as for the wandering ants, the situation is similar to that of more direct emigrations.

The outdegree distribution of the network allows for quantification of the difference in recruitment effort contributed by primary and secondary ants (Figure 4c). For both categories, the outdegree distribution is right skewed, but more so for primary ants than secondary ants. Primary ants have larger mean and variance of outgoing edge numbers, with a few ants exceeding 20 recruitment events (mean ± SD of 2.85 ± 4.03, 5.97 ± 6.78, 3.84 ± 5.03 for colonies 6, 208, and 3004, respectively). Secondary ants nearly always have fewer than five recruitment events and often go as low as not recruiting at all (mean ± SD of 0.44 ± 1.28, 1.12 ± 2.17, 0.9 ± 3.33). Whereas primary ants are often involved in recruitment behaviour across treatments (probability of at least one recruitment of 0.61, 0.76, and 0.65), secondary ants participate more sporadically (probability of 0.14, 0.38, and 0.21).

Additional evidence for the central role played by primary ants is provided by the aggregate recruitment networks that combine all of the emigrations by each colony (Figure 5a and Figure S11). We computed its core and periphery [38] to identify ants that frequently recruit each other (see Figure 5b and Figure S12). The core of a complex network has been shown to have strong connections with the controllability of the underlying networked system [40] and, when its magnitude is large, it is believed to allow for increased flexibility and adaptability [43]. In our experiments, the core is composed of approximately a third of the colony (40%, 23.5%, and 34.4%) most of them primary ants (80%, 73.7%, and 81.8%) and a remaining smaller portion of secondary ants (see Figure 5c and Figure S13); the periphery instead contains at most a small portion of primary ants (4.2%, 8.0%, 0%) and a larger portion of secondary ants (14.6%, 12.9%, 27.3%). For all colonies, we found that the core is a strongly connected network (*i.e.*, it is possible to go from any node to any other node by walking along directed edges). Every ant belonging to the core is therefore recruited at least once across different treatments, highlighting the homogeneous structure of the social network represented by the core.

**Figure 5.**
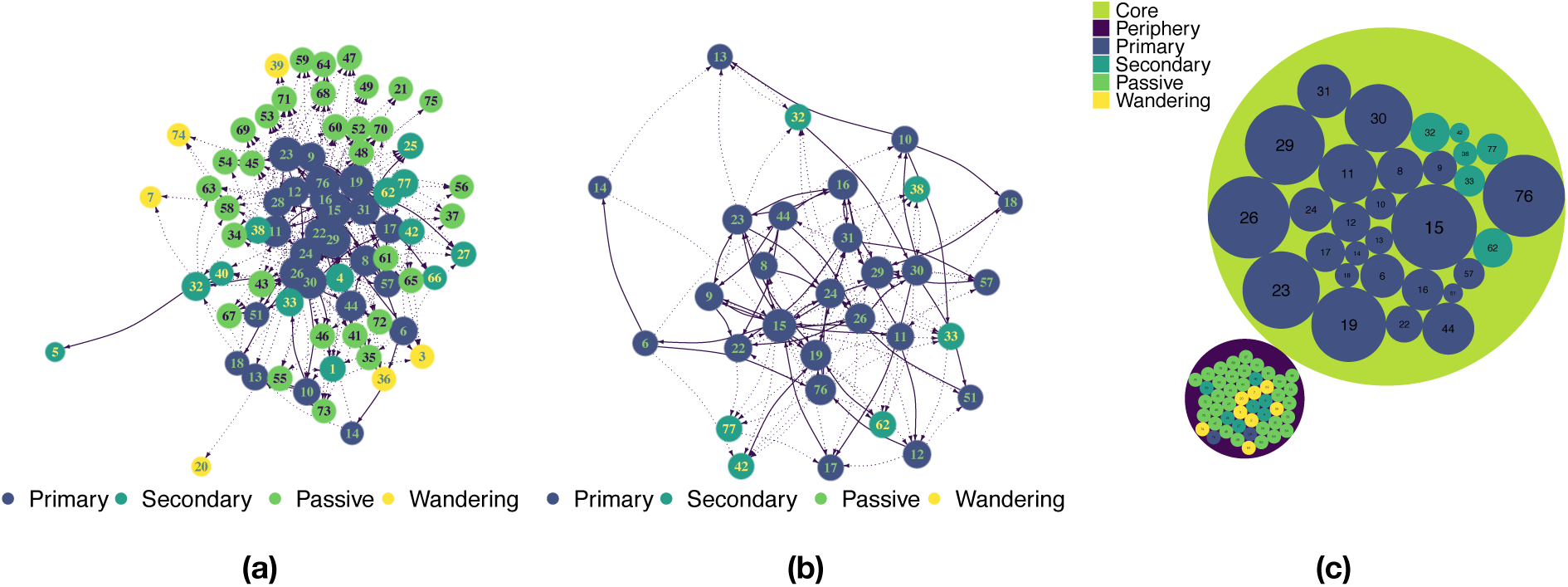
Illustration of aggregate recruitment networks for colony 6. Panel (a) shows the aggregate recruitment network of all treatments for colony 6. Panel (b) shows the core of the network in panel (a). Solid arrows represent tandem runs (both direct and reverse), dotted arrows represent transport events. Isolated nodes are not shown. Panel (c) illustrates the relations between the role covered by each ant as defined from the clustering analysis and their location in the aggregate network (i.e., core or periphery).

### Division of labour

We investigated division of labour during colony emigrations using information-theoretic measures [31]. We considered six different recruitment-related tasks and analysed both worker specialization (*i.e.*, whether workers focus on only a few tasks) and task segregation (i.e., whether certain tasks tend to be performed by the same workers). When considering each worker separately, we found evidence of task specialization across colonies (*DOL*_*indiv*_ = 0.42, *se* = 0.07) but low task segregation (*DOL*_*task*_ = 0.13, *se* = 0.02) which results in a relatively low division of labour (*DOL* = 0.23, *se* = 0.01). When pooling workers into behavioural castes instead, the level of caste specialization is lower (*DOL*_*indiv*_ = 0.28, *se* = 0.07) but that of task segregation among castes is higher (*DOL*_*task*_ = 0.35, *se* = 0.04) resulting in an overall higher division of labor (*DOL* = 0.31, *se* = 0.04). Thus, while the examined tasks are not uniquely performed by specialists, we found evidence that individual workers concentrate more on certain tasks than others and that the problem of finding a new home for the colony is tackled by an organized structure of behavioural castes.

### Spread of information

During emigrations, most workers received information about a potential nest site from other workers through a recruitment event. Only a small proportion of ants gathered this information first-hand and spontaneously initiated recruitment to a new nest (mean ± SD of 11.38% ± 3.28, 13.93% ± 4.23, and 6.96% ± 1.37, respectively, for colonies 6, 208, and 3004). For colony 6, all spontaneous recruiters were primary ants. In colonies 208 and 3004, most were primary ants (mean ± SD of 81.77% ± 13.77 and 93.33% ± 14.9, respectively,) while the remainder were secondary. Primary ants are therefore the initiators of the spread of information about a potential nest. Spreading information, however, is not sufficient to generate an information cascade across all members of the colony. This requires recruited ants to themselves begin recruiting, which depends on the quality of the site being advertised. Indeed, with the exception of one emigration in which recruitment was exclusively focused on the mediocre nest (colony 3004, treatment 3), information about the mediocre nest never cascaded across the members of the colony. In other words, no ant recruited to the mediocre nest ever began recruiting herself to that nest. In contrast, all ants received information about the good nest within two hops of a spontaneous recruiter in the network.

The final choice of the colony seems therefore to depend mainly on the actions of primary workers with secondary workers instead involved in implementing the decision and spreading information about it. Passive and wandering workers are subject to the choice made by active ants. Indeed, whereas both tandem runs and transport are recruitment mechanisms, forward tandem runs (largely performed by primary workers) are more effective in recruiting recruiters than transport events (the primary task of secondary workers) (Table S2). Furthermore, we found ten occurrences in which an ant that was transporting to the mediocre nest switched her efforts to the good nest after being transported there by a nestmate. These ten occurrences happened in the two treatments in which colony 208 was initially split between the good and the mediocre nest (six in treatment 2, four in treatment 3) and involved almost exclusively primary ants (only one ant in treatment 3 was a secondary ant). Moreover, only three out of ten of these events involved ants that were already aware of the location of both nests. This use of transport instead of tandem running to recruit a future recruiter is rather unusual as transports are believed to not allow the transportee to learn the route between two locations [44].

## Discussion

Our analysis confirms previous evidence that colony emigrations by *Temnothorax* ants are organized by an influential minority of workers [18,27–29,45]. This minority, which we call primary ants, makes up about a third of the colony and consists of those workers that participate in tandem running. Primary ants play a critical role in the final choice of the colony, corresponding to that of the decision-making ‘oligarchy’ found in *T. albipennis* [27]. Unlike *T. albipennis*, we found no specialization on specific tandem run roles; that is, there was no negative correlation between participation as a leader and participation as a follower.

In addition to primary ants, we identify three other behavioural castes. About a fifth of the workforce consists of secondary ants that are generally transported to a candidate nest by a primary ant and soon begin to carry nestmates there from the old nest. Although secondary ants contribute to the implementation of the colony’s choice, they do not seem to contribute directly to the decision-making process. However, they might indirectly influence the choice by hastening achievement of the quorum sensed by primary ants [18]. Nearly all remaining workers (*i.e.*, about half of the colony) consist of passive ants – workers that participate in the emigration only by being carried to a candidate nest. Passive ants likely correspond to the lazy ants described in studies of task allocation [42]. We also found a small proportion of workers, 3–12% of the colony, that do not fit any of the other castes. While these wandering workers do not seem to participate in decision-making or implementation, they still perform an unusually high number of visits to candidate nests. We speculate that they are foragers in search of food.

The different behavioural castes fill distinct roles in processing the information needed for nest site choice. Primary ants, with their reliable use of tandem runs, discover potential nest sites and then ensure that the newly gathered information spreads through the colony. They also process this information by comparing different options both individually, when they visit both nests before they start recruiting, and collectively, by modulating recruitment efforts as a function of quality [34]. Among primary ants, only about 10% of the colony actually gather this information first-hand as a result of random exploration of the environment. Tandem run recruitment of other active ants (predominantly primary but likely including a minority of secondary ants) slowly processes the gathered information, resulting in the buildup of a quorum in one or (occasionally) more sites. As evidenced by two treatments of colony 208, the initial recruitment leading to the establishment of a quorum can sometimes rely on transport of adult workers between competing nests in a way that is reminiscent of cross-inhibition in honeybees [46]. Once quorum is reached, transport becomes the recruitment mechanism of choice and the remaining members of the colony are carried to the chosen site. During this phase, secondary ants assist the implementation of the minority’s collective decision; by transporting other secondary ants as well as passive ants, they spread information concerning the colony decision to the rest of the society. Passive ants might also contribute indirectly to the establishment of a quorum by increasing encounter rates at a site to which they are carried [18]. The possible contribution of wandering ants still remains unclear.

Although the behaviour of workers (and therefore their membership in a particular behavioural caste) was stable over the course of five emigrations, we know that individual workers change the tasks they perform as a function of their developmental age [23]. Furthermore, even though the proportion of primary ants seems relatively stable [18,27,28], it remains to be shown that this is still true for each behavioural caste we identified. Differences in these proportions might affect the information-processing ability of the collective and result in different colony personalities manifested as different tradeoffs between speed and accuracy [47].

Division of labour is often credited with the ecological success of social insects [48]. In social animals more generally, there is growing evidence for major impacts of behavioural variation on group behaviour and fitness [49,50]. An open question is whether the behavioural distinctions documented here offer any functional advantages to the colony. Of particular interest is the difference between primary and secondary ants. Even if we acknowledge that this distinction is a coarse-grained simplification of more continuous variation, our results show that the ants most directly involved in the emigration vary strongly and consistently in behaviour. Future work can determine whether dividing labour in this way enhances the quality of collective decision-making.

## Acknowledgments

This work was partially supported by NSF grant No. PHY-1505048. We are grateful to many undergraduate students who aided in video analysis, including Jeni Briner, Courtney Bruce, Savannah Christy, Sam Fox, Aaron Houglum, Glenn LeSueur, Nick Poetsch, Shay Richardson, and Meghan Stewart.

