## Supplementary information file for "Division of labour promotes the spread of information in colony emigrations by the ant *Temnothorax rugatulus*"

### 1. Clustering analysis

#### 1.1 High level clustering

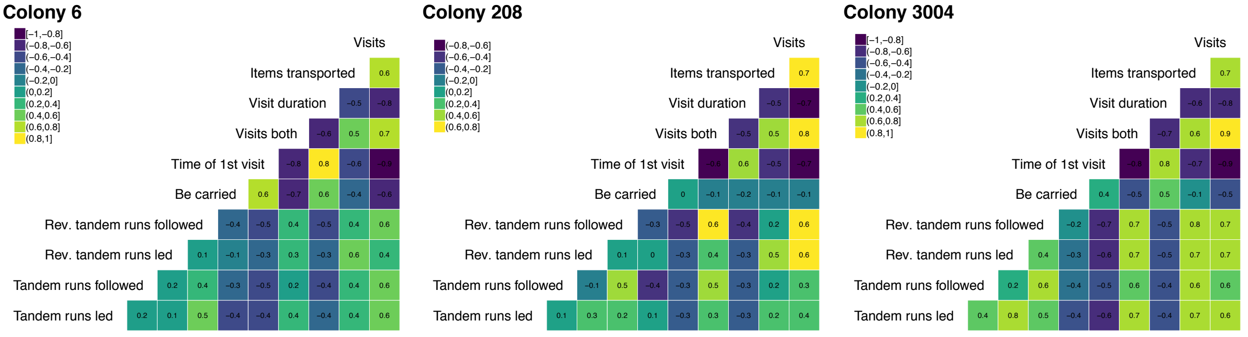


(a) (b) (c)

*Figure S1 Matrix of Pearson correlation coefficients between all behavioural features considered at the higher level of the hierarchy for (a) colony 6, (b) colony 208, and (c) colony 3004.*

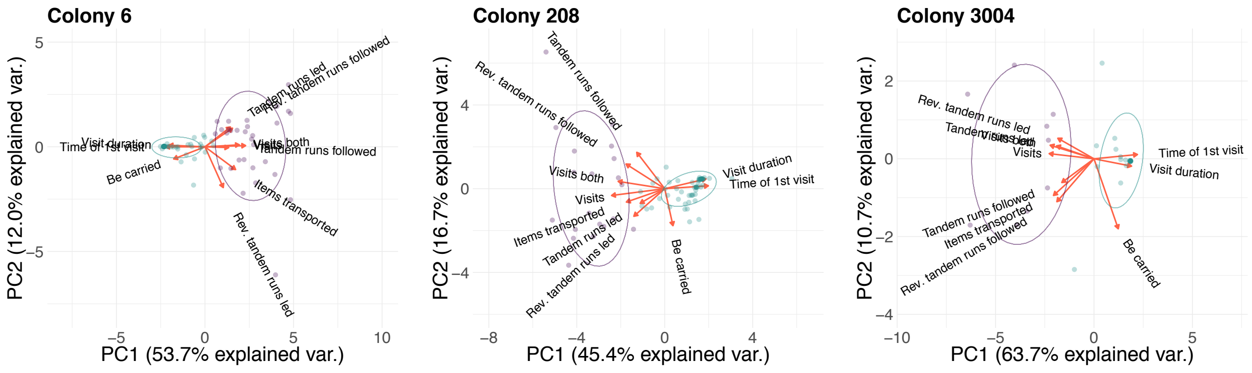


(a) (b) (c)

*Figure S2 PCA representation over the first two components of the loadings for the behavioural features considered at the higher level of the hierarchy for (a) colony 6, (b) colony 208, and (c) colony 3004. Coloured points represent clustered workers (purple colour for active workers, green colour for inactive workers).*

### 1.2 Low level clustering

In the case of active workers, we considered all behavioural features included at the higher level of the hierarchy plus three additional features related to transport: the average duration of a transport event, the time of first transport, and the cumulative visit duration before transporting an item. Behavioural features generally had a low level of pairwise correlation with the exception of two sets of features with high intra-set correlation (Pearson correlation coefficients in Figure S3). One set included features related to transport events, and the other included features related to the timing of visits. As the PCA (see Figure S4) showed two separate groups of workers, we further divided active workers in two groups: *primary* workers and *secondary* workers.

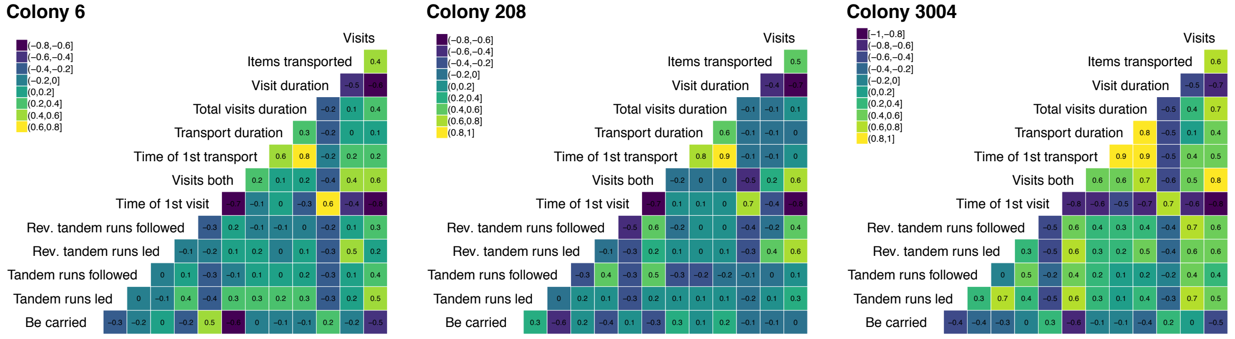


(a) (b) (c)

*Figure S3 Matrix of Pearson correlation coefficients between all behavioural features considered at the lower level of the hierarchy for active workers of (a) colony 6, (b) colony 208, and (c) colony 3004.*

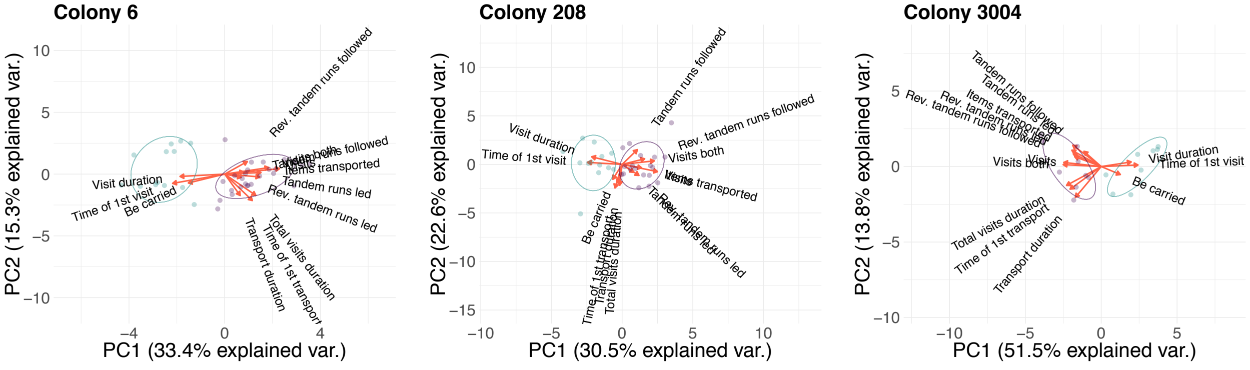


(a) (b) (c)

*Figure S4 PCA representation over the first two components of the loadings for the behavioural features considered at the lower level of the hierarchy for active workers of (a) colony 6, (b) colony 208, and (c) colony 3004. Coloured points represent clustered workers (purple colour for primary workers, green colour for secondary workers).*

In the case of inactive workers, we considered only a few behavioural features: the number of times they have been carried to a nest, the number of visits, their average duration, time of the first visit, and whether they visited both nests (this feature was not considered for colony 3004 as inactive workers never visited both nests). The number of visits to a candidate nest is positively correlated to the likelihood of visiting both nests whereas the mean visit duration is positively correlated with the number of times a worker is carried to a nest (Figure S5). From the PCA (Figure S6), we observe that loadings are still divided in two separate sets pointing in opposite directions. We therefore clustered inactive workers in two groups: *passive* workers and *wandering* workers.

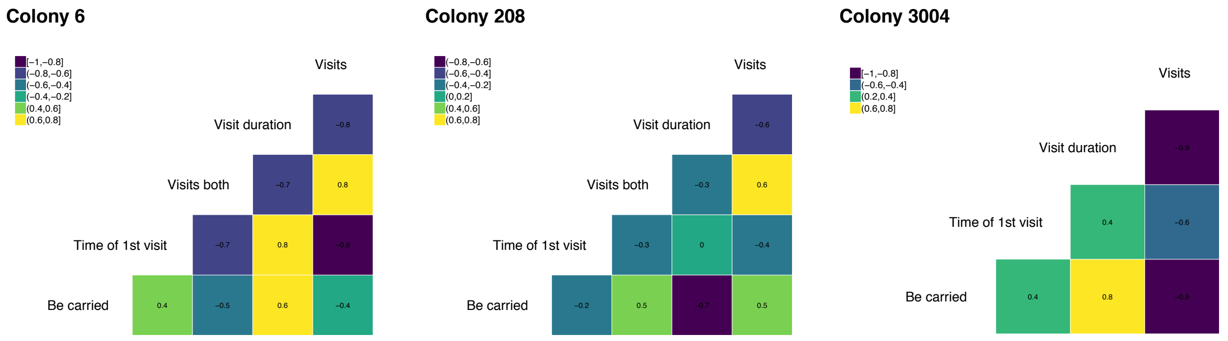


(a) (b) (c)

*Figure S5 Matrix of Pearson correlation coefficients between all behavioural features considered at the lower level of the hierarchy for inactive workers of (a) colony 6, (b) colony 208, and (c) colony 3004.*

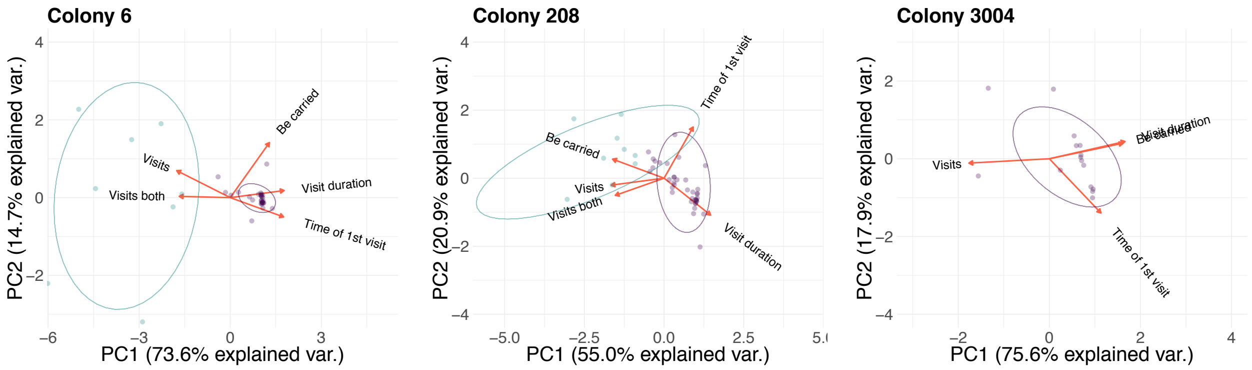


(a) (b) (c)

*Figure S6 PCA representation over the first two components of the loadings for the behavioural features considered at the lower level of the hierarchy for inactive workers of (a) colony 6, (b) colony 208, and (c) colony 3004. Coloured points represent clustered workers (purple colour for passive workers, green colour for wandering workers).*

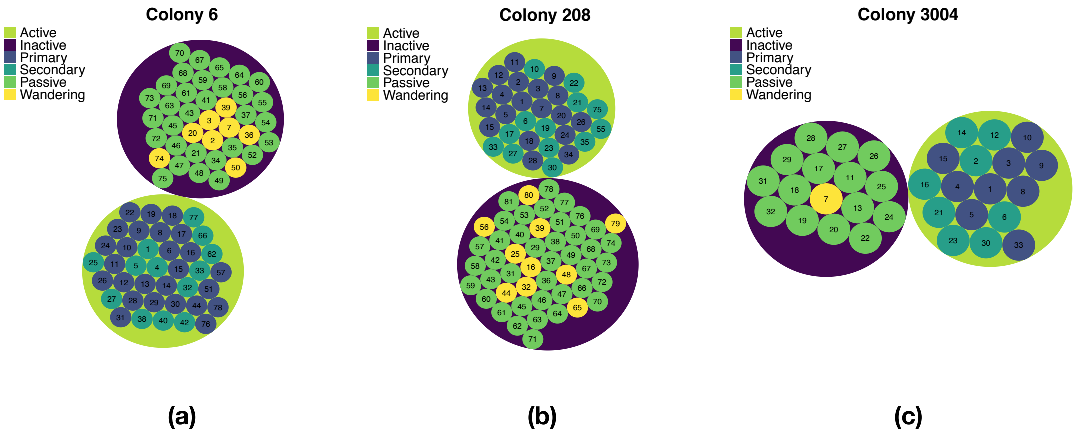


*Figure S7 Circle plots representing the overall results of the clustering of ants for (a) colony 6, (b) colony 208, and (c) colony 3004.*

### 2. Network analysis

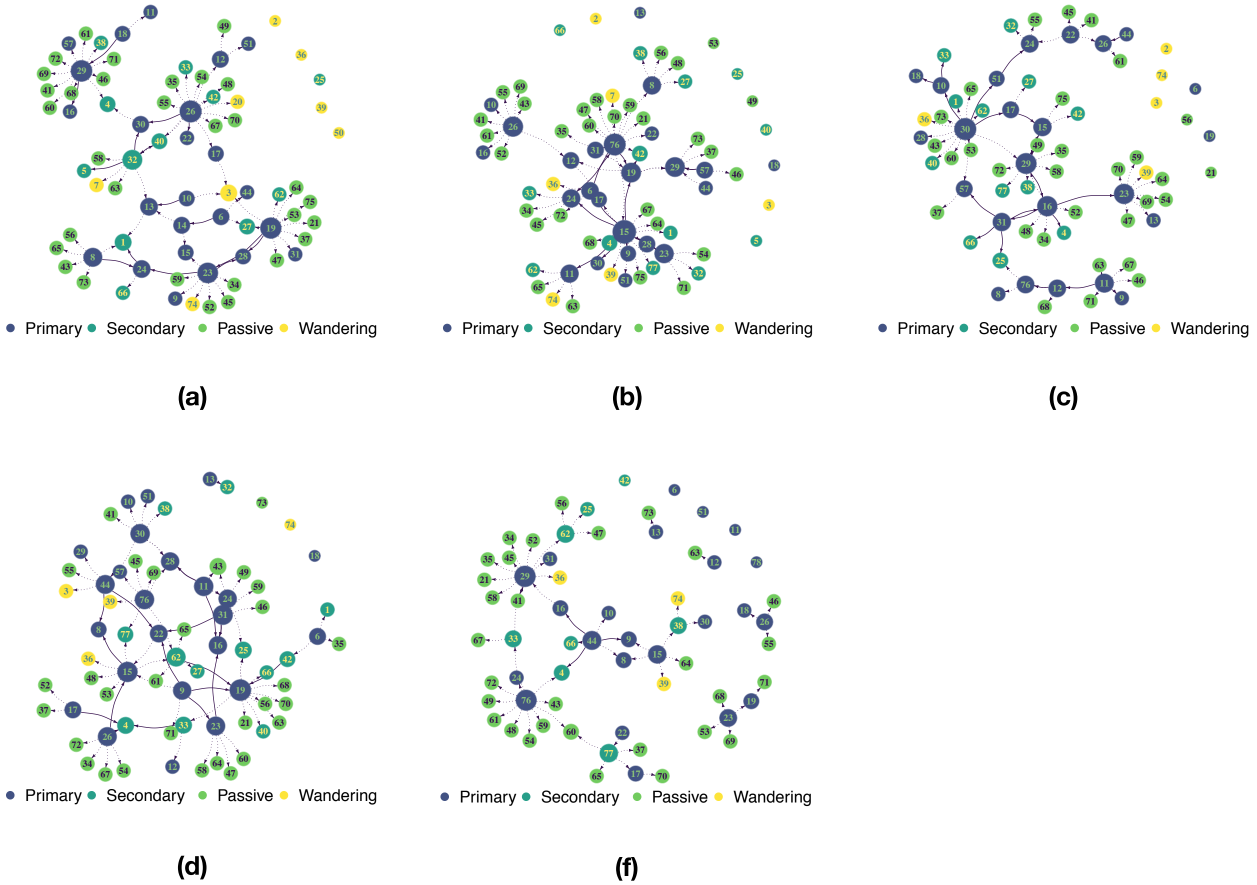


*Figure S8 Illustration of recruitment networks for treatment 1–5 of colony 6. Solid arrows represent tandem runs (both forward and reverse), dotted arrows represent transport events.*

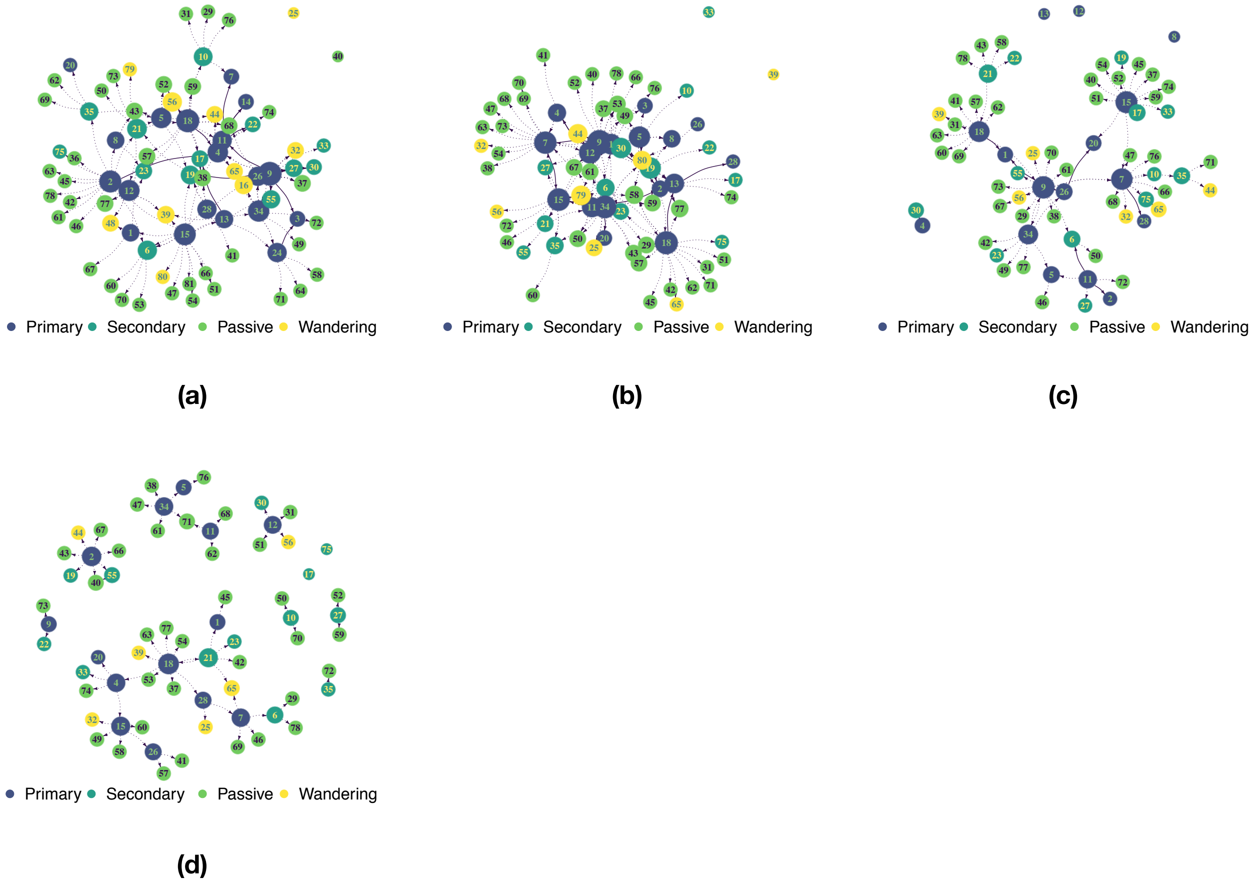


*Figure S9 Illustration of recruitment networks for treatment 2–5 of colony 208. Solid arrows represent tandem runs (both forward and reverse), dotted arrows represent transport events.*

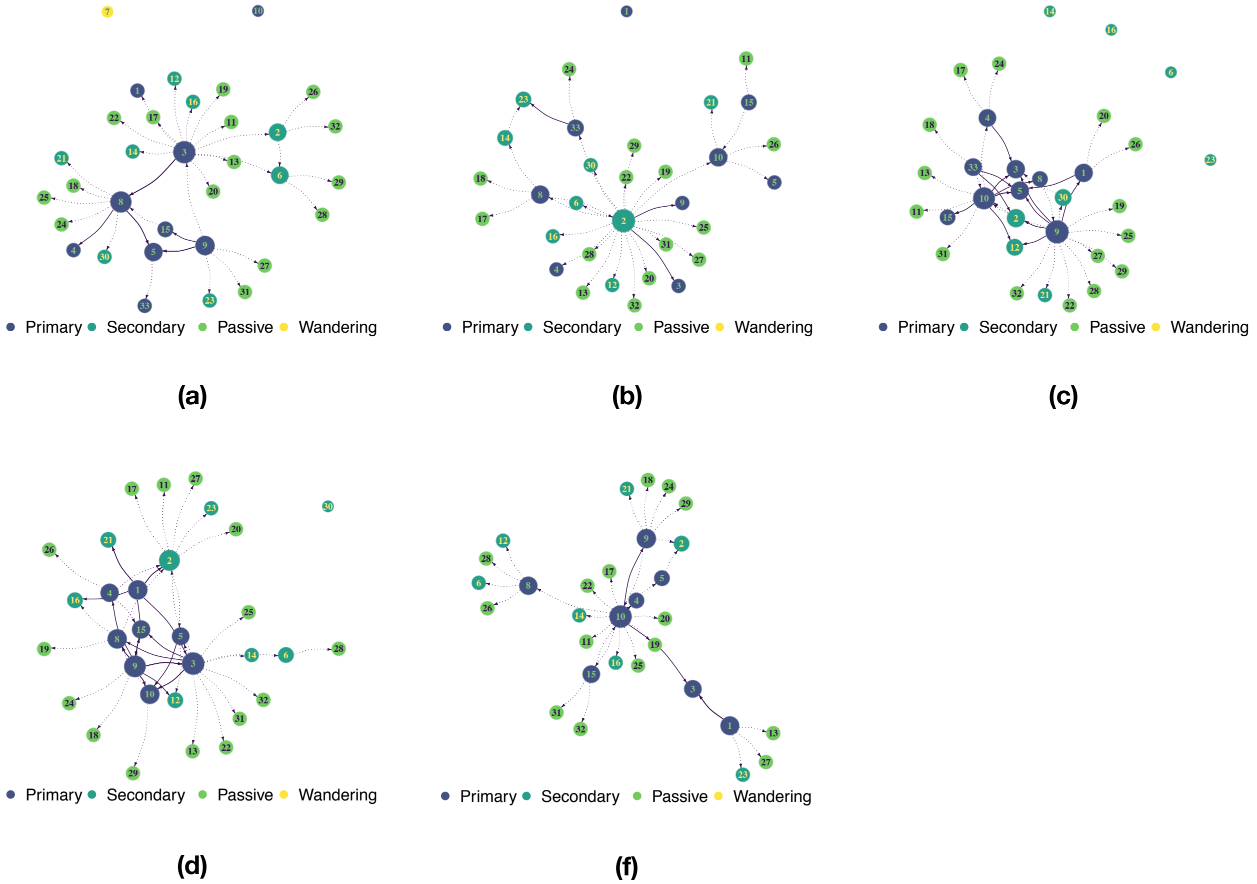


*Figure S10 Illustration of recruitment networks for treatment 1–5 of colony 3004. Solid arrows represent tandem runs (both forward and reverse), dotted arrows represent transport events.*

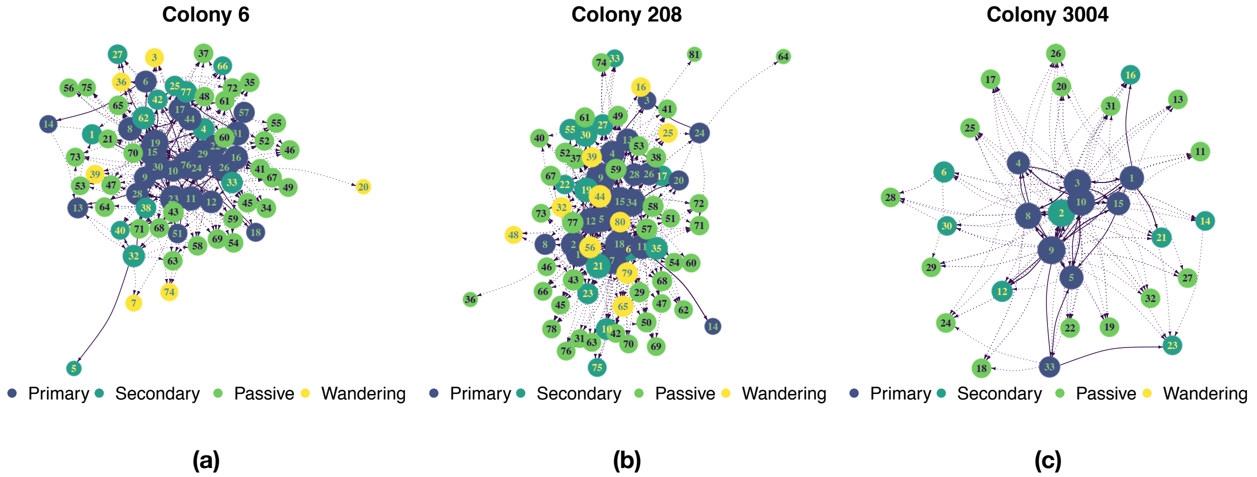


*Figure S11 Aggregate recruitment network for (a) colony 6, (b) colony 208, and (c) colony 3004. Solid arrows represent tandem runs (both forward and reverse), dotted arrows represent transport events. Isolated nodes are not shown.*

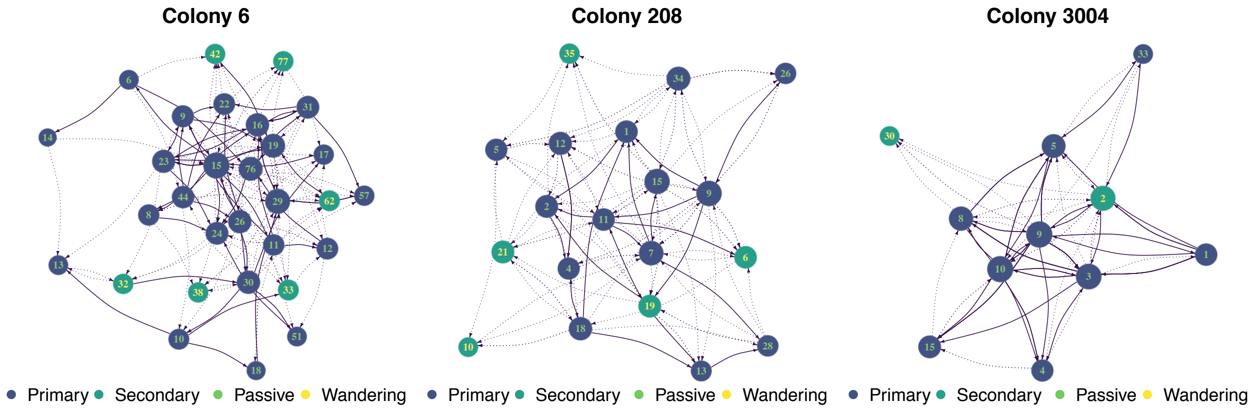


(a) (b) (c)

*Figure S12 Core of the aggregate recruitment network for (a) colony 6, (b) colony 208, and (c) colony 3004. Solid arrows represent tandem runs (both direct and reverse), dotted arrows represent transport events. Isolated nodes not shown.*

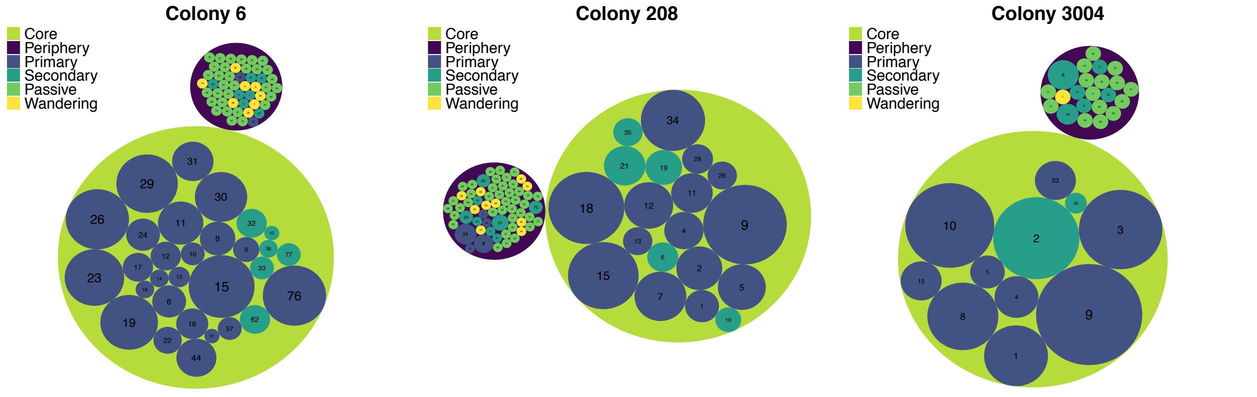


(a) (b) (c)

*Figure S13 Illustration of the relations between the role covered by each ant as defined from the clustering analysis and their location in the aggregate network (i.e., core versus periphery), respectively, (a) for colony 6, (b) for colony 208, and (c) for colony 3004.*

*Table S1 Measures of division of labour for each colony and their mean and standard deviation divided, respectively, in ant versus task and role versus task.*

| | (ant, task) | | | *(role, task)* | | |
|---|---|---|---|---|---|---|
| *Colony* | $DOI$ | $DOI_{indiv}$ | $DOI_{task}$ | $DOI$ | $DOI_{indiv}$ | $DOI_{task}$ |
| 6 | *0.23* | *0.43* | *0.12* | *0.33* | *0.28* | *0.39* |
| 208 | *0.24* | *0.48* | *0.12* | *0.33* | *0.34* | *0.32* |
| 3004 | *0.23* | *0.34* | *0.15* | *0.26* | *0.21* | *0.33* |
| $mean \pm sd$ | $0.23 \pm 0.01$ | $0.42 \pm 0.07$ | $0.13 \pm 0.02$ | $0.31 \pm 0.04$ | $0.28 \pm 0.07$ | $0.35 \pm 0.04$ |

*Table S2 Number and proportion of recruitment events that turned the recruited worker into a recruiter herself. Results divided for forward tandem runs and transport events.*

| *Colony* | *Forward tandem runs* | *Transports of adults* |
|---|---|---|
| 6 | $13/23 = 0.57$ | $19/257 = 0.07$ |
| 208 | $3/9 = 0.33$ | $31/240 = 0.13$ |
| 3004 | $8/15 = 0.53$ | $14/121 = 0.12$ |
